# Fire shifts the structure and dynamics of woody plant communities in a mediterranean coastal dune ecosystem

**DOI:** 10.1101/2025.01.14.633058

**Authors:** André F. Mira, Nagore G. Medina, Sergio Chozas, Raquel Divieso, Juan Castro-Rivadeneyra, Mari Cruz Díaz-Barradas, Joaquín Hortal

## Abstract

Coastal dune ecosystems are dynamic and fragile environments shaped by complex interactions between vegetation and its characteristic abiotic factors. This study investigates the effects of the wildfire that affected the Doñana Nacional Park (SW Spain) in 2017 on the composition and structure of woody plant communities in two adjacent Mediterranean coastal dune areas: one affected by fire (Cuesta Maneli) and one unburned (Laguna del Jaral). Using a combination of several analyses (such as non-metric multidimensional scaling (NMDS); Mantel tests; and co-occurrence analyses), we examined how fire influences species distribution, abundance, and their interactions along a coastal-inland gradient. Our results reveal that fire homogenizes woody plant community composition across the coastal-inland gradient, reducing spatial differentiation and increasing overall species abundance in the burned area. Soil characteristics influenced species’ composition significantly only in the unburned site, suggesting that fire disrupted abiotic filtering processes. Co-occurrence patterns indicate more neutral interactions in the burned area, consistent with reduced competition and increased resource availability post-disturbance. In contrast, the unburned site displayed structured communities with stronger negative co-occurrence at the smaller scales and spatial clustering of species. Spatial pattern analyses showed that positive co-occurrence was more prevalent in the burned area, particularly at smaller spatial scales, indicating potential facilitation effects during early post-fire recovery. These findings highlight how fire reshapes community structure and dynamics after succession. Understanding these processes is critical for effective conservation and management of Mediterranean coastal dune ecosystems facing increasing anthropogenic pressures and climate change-related disturbances.

## Introduction

Coastal dune ecosystems are considered unique as they represent the transitional belt between terrestrial and marine environments ^1,2^. They accommodate a distinctive combination of characteristics such as high temperature variability, low soil fertility, limited water availability, varied geomorphology, sand movement, strong winds, and salt spray that makes dune ecosystems often highly patchy and dynamic ^3–5^. The combination of such elements creates a scenario in which the vegetation present in these systems faces not only the adversities imposed by the overall environment but also the patch effects created by the heterogeneous distribution of stresses present at the finer spatial scales ^6^. These dynamics generate reciprocal interactions between vegetation and spatial and abiotic characteristics, as plants stabilize and promote the conditions most favorable to them ^4^. Such feedback between vegetation and the environment promotes a locally heterogeneous, patchy plant community ^7^. This often results in increased habitat complexity further inland due to greater vegetation cover, forming a landward stability gradient, with increased vegetation helping to mitigate physical stressors ^3^. Furthermore, this interplay establishes areas of highly specific assemblages (or stability domains), since dune-modifier plants are able to establish conditions that promote/demote the presence of species from the regional pool species ^4,8^. Thus, to achieve a comprehensive understanding of these communities, it is essential to consider the mechanisms that filter species to the local scale, as well as how interactions within the local community influence the broader regional species pool ^9^.

Moreover, these intrinsic characteristics also render them fragile ecosystems as they are vulnerable to numerous potential threats. Indeed, coastal dune ecosystems are recognized by the *International Union for Conservation of Nature* as a highly vulnerable ecosystem as they are currently very susceptible to massive biodiversity loss ^10–12^. In fact, these ecosystems accommodate unique phytocoenosis that encompass a high proportion of endemic species and harbor habitats that are vulnerable to both natural and human perturbations ^5^. Mediterranean coastal dunes are a prime example, as they present a profuse number of unique habitats (with international conservation interests ^13^) that contain highly specialized flora and fauna ^2^. However, factors such as the expansion of agriculture, afforestation, unconstrained tourism, and the enlargement of urban areas, have been steadily increasing the vulnerability of this ecosystem in the recent decades ^14,15^. As these anthropogenic activities increase in number and area, they facilitate many indirect processes which impose a threat to natural biodiversity ^16,17^.

One of the most menacing perturbations to coastal environments is fire, which can arise and be augmented by both anthropogenic and natural factors. For the Mediterranean regions in particular, the last decades have brought a significant increase in the frequency of fires and total area burned ^18,19^. As fire frequency and intensity increase, these factors have been identified as key drivers that not only influence vegetation dynamics but can also lead to shifts in species composition and the structure of plant communities ^20,21^. Furthermore, fire is also known for being a factor that triggers erosion and sand loss, especially when combined with extreme climatic conditions ^22,23^. Such events are also amplified by the amount of vegetation that burns, which decreases the stability of dunes leading to more sand particles being blown away ^22,24,25^. Considering the potential effects of fire perturbation on coastal Mediterranean ecosystems, the regeneration of the plant communities may not follow the expected successional pattern, as certain shrub species may fail to recover and/or be replaced by others ^26^. Although the impact of fire in coastal Mediterranean ecosystems has been extensively studied ^27–30^, there is still limited understanding of how changes in species composition, driven by fire perturbation, alter the interactions among species within the broader community.

Since Mediterranean coastal dunes are exposed to perturbations that can significantly change their core dynamics, it is imperative to gather knowledge of the relationships between local plant communities and the environmental conditions they reside in to achieve effective conservation planning. Coastal dunes plant community structure and diversity is the outcome of the interplay between mechanisms that occur at larger scales (both abiotic and biotic) and at the local scale (environmental stress and species interactions ^31–33^). At larger scales, biogeographic processes shape plant species distribution patterns both within and across regions. Species diversity across these broader scales is often influenced by regional climatic gradients, which play a key role in determining how species are distributed and interact over extensive areas ^34,35^. On the other hand, at the local scales plant communities face other challenges. For instance, in the abiotic component, local communities are shaped by intricate environment gradients generated by the combination of several factors such as salt spray, substrate mobility, composition, and nutrients ^7,36–38^. Furthermore, biotic factors also interpose a filter at the local scale that affects not only the plant community but also how some species react to certain abiotic conditions ^6,7^. Several studies have explored the relationship between positive and negative species interactions along gradients of dune environments ^39,40^. However, there is still limited research on how interactions, whether positive or negative, vary with the distance between individuals within specific spatial segments of the inland-coastal gradient, and how these dynamics may be altered by fire disturbances.

Given this context, our objective was to investigate the composition and structure of woody plant communities in two adjacent Mediterranean coastal dune areas in the SW of Spain: one affected by a fire in 2017 and the other unburned. We aim to understand how fire can alter the succession and dynamics within these already fragile communities. By burning biomass, fire releases nutrients into the soil which can lower competition, further enhancing opportunities for plant establishment and growth. Thus, for community composition we anticipate that the burned area will harbor a higher number of individuals and species, as the fire-induced vegetation gaps are likely to facilitate colonization. Furthermore, we hypothesize that fire will homogenize the composition of the coastal dune plant community along the coastal-inland gradient, resulting in decreased variation in species distribution across this gradient. Regarding the structure of the community, we expect to find more random species associations in the burned area, reflecting a less structured and more opportunistic assembly of species following the succession post-fire disturbance. In contrast, the unburned area is expected to exhibit a more spatially structured community, characterized by well-established species interactions that maintain a more predictable and organized pattern of species distribution across the landscape.

## Materials and Methods

### Study region

The study region is located within the coastal dune system of the Doñana Natural Park, southwest (SW) of Spain (Fig. 1). This coastal system is composed of an extensive area, encompassing various geomorphological features such as young moving dunes, fossil dunes, eolian sand sheets, and a beach, extending along approximately 30 kilometers of the Atlantic coast with 1 kilometer width ^41,42^. As of today, the study region contains an environmental boundary delineated by the 2017 fire’s extent, which impacted the western (W) portion of the coastal dune system. Before the fire, the vegetation of this coastal region was composed of an extensive pinewood of mostly *Pinus pinea L*. (esulting from reforestation efforts in the 1950s) and some scrub species ^41–44^. After the fire, resprouter shrub species such as *Osyris lanceolata* (Hochst. & Steud) and *Corema album* (D. Don) started recolonizing the landscape, due to their regenerative growth capabilities. Afterwards, the spring rains of 2018 significantly enhanced the recolonization of pyrophytic seeder species. In particular, *Halimium halimifolium* ((L.) Willk.), *Cistus salviifolius* (L.), and various herbaceous annuals proliferated during this period, thereby contributing to the development of the plant community. Over time, a distinct vegetation community structure has emerged in the western area, while the eastern (E) area remains unchanged, since it was not affected by the fire.

**Figure 1 –.**
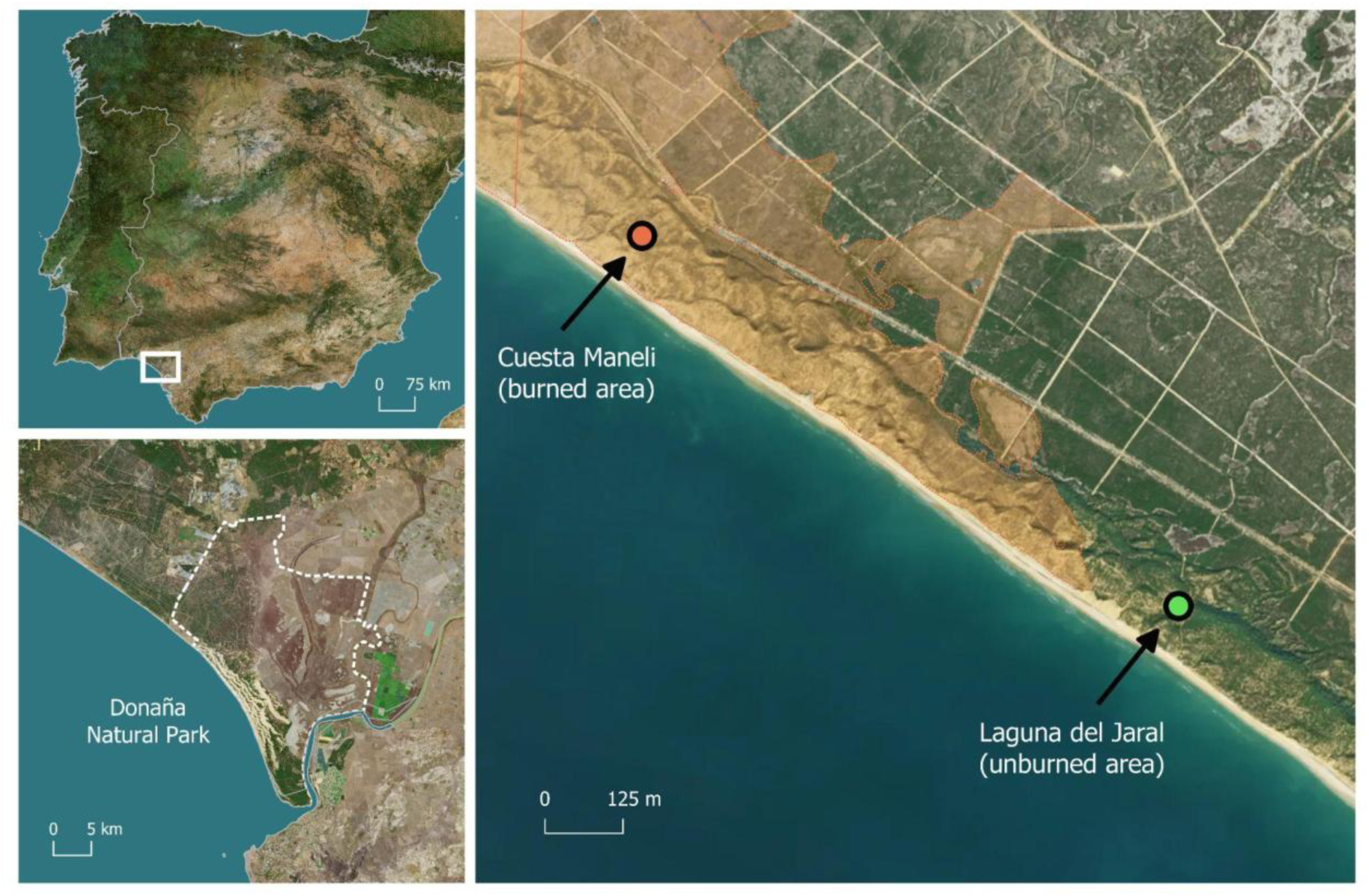
Location of both sampling areas in the SW of Spain.

### Sampling methodology

Sampling was conducted in two areas: Cuesta Maneli in the west, affected by the fire (referred to as the “burned area”), and Laguna del Jaral in the east, which remained unaffected (the “unburned area”). These areas are 6.5 kilometers apart. Overall, three 200 meters transects (per study area) were drawn along a coast-inland gradient characterizing three distinct zones: a coastal zone located 200 meters from the shoreline, an intermediate zone 500 meters inland, and an interior zone 850 meters from the shore. Altogether, 36 sampling plots were established (6 per transect). Each plot comprised a 2-by-2-meter square divided into 16 sub-plots, within which woody vegetation species inventories were carried out. Furthermore, for each sampling plot, composed soil samples were collected for soil element analysis. Two sampling seasons were carried out: the burned area in January 2022 and the unburned in January 2023.

### Soil Analysis

The obtained composite soil samples were air-dried and passed through a 2-mm sieve before laboratory analysis. For each prepared samples, the following test were performed: a) *pH measurement*: pH was determined using water (pH H₂O) and potassium chloride (pH KCl) extraction; b) *Organic Matter*: Quantified as a percentage of the total sample; c) *Nitrogen Content*: Total nitrogen measured using the Kjeldahl method; d) *Available Phosphorus and Potassium*: Analyzed to determine the concentrations accessible to plants; e) *Exchangeable Potassium, Calcium, and Magnesium*: Measured to assess the nutrient availability; f) *Carbonates (Bernard Method)*: Carbonate content was determined; g) *Soil Texture (Hydrometer Method)*: Analyzed using a densimeter; h) *Active Lime*: Determined to assess the presence of active calcium carbonate; i) *Nitrate and Ammonium*: Extracted using KCl and analyzed; j) *Total Nutrients and Micronutrients*: Assessed using ICP-OES and ICP-MS for a broad spectrum of elements; k) *Available Micronutrients (EDTA Extraction)*: Analyzed using ICP-OES; l) Electrical Conductivity (EC).

### Statistical Analysis

All statistical analysis was performed using RStudio version 4.3.1^45^.

### Woody plant species community composition

To understand how the composition of woody plant species community changes within and between the sampled areas, several analyses were performed. Firstly, to study the effect of spatial distance and soil parameters on the composition of woody species assemblages we used a Multivariate Mantel test ^46,47^, using the *‘mantel’* function of the *‘vegan’* R package ^48^. To do so, three data matrices were created for each sampling area: 1) Abundance Matrix: containing Bray-Curtis dissimilarity indices computed for woody plant abundance; 2) Geographical Matrix: Longitude and latitude data were extracted and converted into distance matrices using the Haversine formula; and 3) Environmental Soil Characteristics Matrix: in which Euclidean distances were calculated from scaled soil characteristics data matrices. The performed Mantel test allowed us to estimate the linear relationship between pairwise combinations of the three standardized matrices.

Then, to explore how the variation in the soil variables changed across the studied plots, a Principal Component Analysis (PCA) was performed. Prior to the analysis, all soil variable data was standardized (mean-centered and scaled to unit variance) to account for differences in measurement scales among the variables. The PCA was conducted using the ‘rda’ function of the *‘*vegan*’* R package ^48^. Additionally, a Non-Metric Multidimensional Scaling (NMDS) analysis was performed to explore the patterns in woody plant species composition across the sampling plots. The analysis was conducted using the ‘MetaMDS’ function from the *‘*vegan*’* R package. Moreover, the previously obtained PCA axes were superimposed onto the NMDS plot to allow a visual inspection of the relationship between the ordination and environmental variables. An environmental fit analysis was performed, using the ‘envfit’ function from the *‘*vegan*’* R package, to determine the relationship between the NMDS ordination space and the PCA axis, as well as categorical factors such as area and transect. The analysis was conducted with 1,000 permutations to assess statistical significance.

Furthermore, an indicator species analysis was conducted using the ‘multipatt’ function from the ‘indicspecies’ package in R ^49^. This approach identified species significantly associated with a specific area or transects, along with their IndVal statistics and p-values.

Finally, to assess the effect of fire and zone on the abundance of woody species individuals, a linear model (‘lm’ function) was employed, and afterwards an analysis of variance (Anova) was conducted to evaluate the significance of the chosen factors. To determine the appropriate model to analyze the richness of woody species, two generalized linear models (‘glm’ function) using site and transect as explanatory variables were fitted: a) using a Gaussian distribution; and b) using a Poisson distribution. The residual plots for both models were examined to assess fit. The Poisson model provided a better fit as indicated by the residual diagnostics and was used in further analysis. An analysis of variance (Anova) was then performed on the Poisson model to determine the significance of the used factors.

### Woody plant species community structure

To study the structure of the woody plant community, based on the spatial organization of the species, we first constructed a presence/absence ‘site x species’ matrix to analyze co-occurrence patterns. Species co-occurrence was analyzed using the ‘EcoSim’ package ^50^. The ‘EcoSim’ package allows us to test patterns in species communities, through the comparison of randomly generated matrices with the observed presence-absence matrices. The “Sim9” algorithm proposed by Gotelli ^51^ was chosen to quantify the mean number of “checkerboard units” among all possible pairs of species in the used matrices. Using this algorithm, a C-score ^52^ is generated for the observed and the simulated presence-absence matrices, which allow us to infer about three spatial distribution scenarios:

a. If the observed matrix C-score is significantly higher than the C-score from the simulated matrix (species co-occur less often than expected by chance) it suggests spatial distributions may be influenced by negative species associations.
b. In the case that the observed matrix C-score is significantly lower (species co-occur more often than expected by chance), it indicates that positive associations between species may influence spatial distributions.
c. If the observed matrix C-score does not differ from the simulated matrix C-score it suggests that species co-occur as expected by chance.

Statistical significance was calculated by generating 30000 random matrices from which the C-score for each matrix was calculated and compared from the real presence-absence data matrix, if the observed matrix C-score was greater than 95% of the highest or less than 95% of the lowest C scores from the randomly generated matrices it was considered significant.

To better understand the structure of the woody plant communities, a co-occurrence network was developed to analyze the spatial organization of woody plant communities across different transects. This was done by applyingthe ratio index of adjacency (using the ‘get_network’ function from the ‘asnipe’ R package ^53^), which gauges the strength of association between species pairs. Afterwards, based on the co-occurrence network, we applied the Louvain algorithm ^54^ to identify modules of interacting species and assess the strength of the community structure. This process was repeated for each area and year combination, and we compared the community structures across different years and areas. The co-occurrence networks were created using the “igraph” package in R ^55^.

Furthermore, spatial point pattern analysis ^56^ was performed to understand how species’ spatial distribution is affected by other species and how their spatial associations change with distance. Firstly, a distance data matrix was created, based on the centroid distance of the sub-plots in which individuals were identified in each sampling plot. In our analysis, the ‘Kcross’ function from the ‘spatstat’ package ^57^ was used to evaluate the relationship between all pairs of species present in each plot. For each pair, the function was computed over a distance range of 0 to 1 meter, with a resolution of 0.001 meters, and Ripley’s correction was applied to account for edge effects. However, since the distance data contained in the data matrix was based on the sub-plot centroid distance, only three distances were considered: ***Distance I*** - 0.001 meters; ***Distance II*** - 0.5 meters; and ***Distance III*** - 1 meter. Afterwards, a data matrix was created for each pair of species with the type of spatial association (attraction or repulsion) for each distance. To understand how *fire* and *transect* affect the type of association, a logistic regression model using the ‘glm’ function was built in R with a binomial family and logit link. All models assumed the burned area and the coastal transect as intercepts. Lastly, to further investigate how fire affected the structure of the woody plant community, we analyzed how the spatial associations (number of positive and negative associations) of the six species that were found in both sampling areas were affected by the fire, by applying logistic regression models using the ‘glm’ function was built in R with a Poisson family and logit link.

## Results

### Drivers of community composition

The Mantel test results identified the main relationships between woody species composition, geographic location, and soil characteristics across sampling sites (Fig. 2). In the burned area, species composition showed a significant positive correlation with geographic distance (Mantel’s R = 0.26, p = 0.006), while no significant correlation with soil characteristics was observed (Mantel’s R = 0.03, p = 0.39). In contrast, in the unburned area, species composition was strongly and positively correlated with both geographic distance (Mantel’s R = 0.50, p = 0.0002) and soil characteristics (Mantel’s R = 0.35, p = 0.004).

**Figure 2 –.**
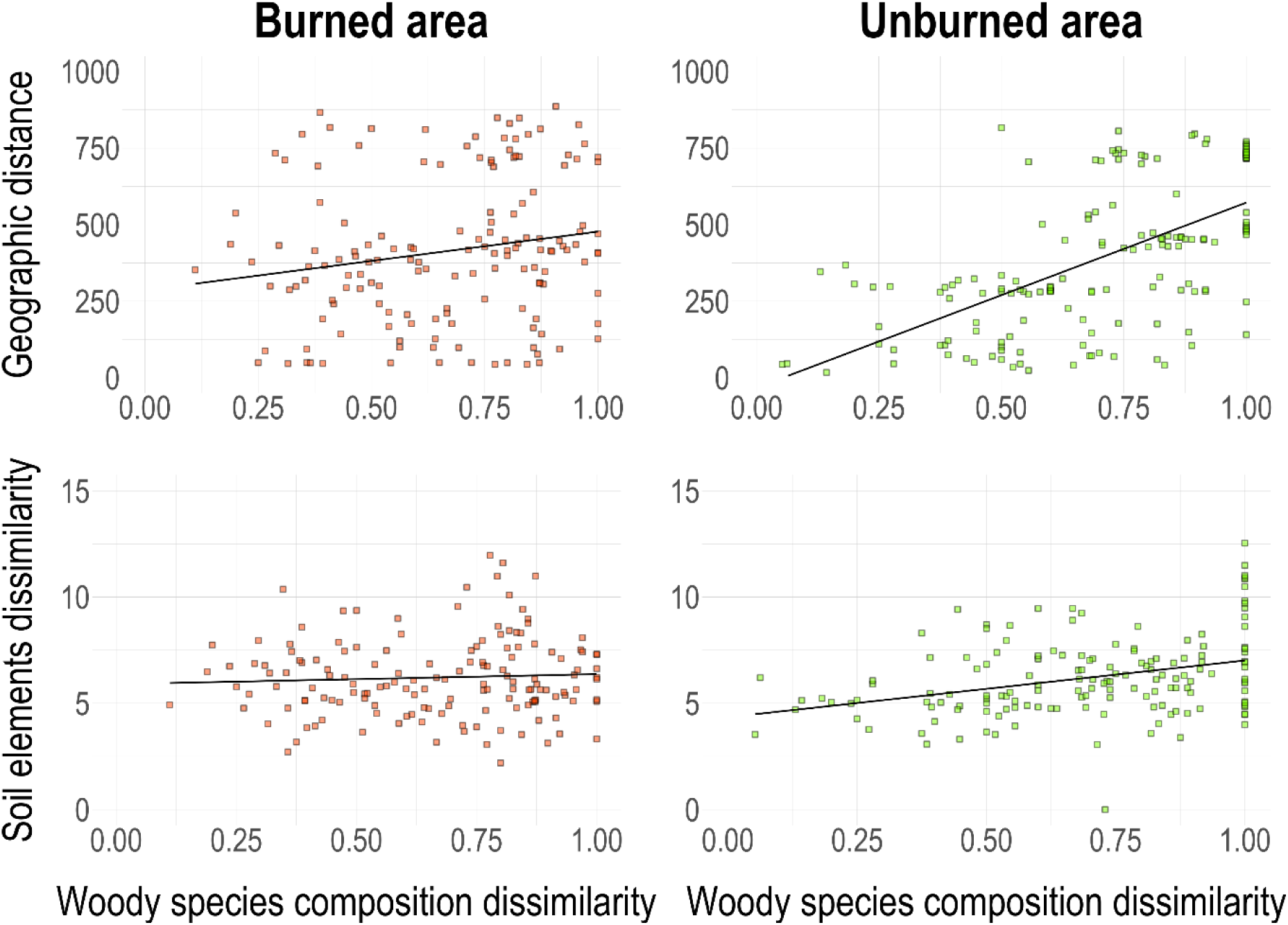
The figure presents the relationship between the woody species composition dissimilarity and the two used variables: geographic distance (top row) and soil elements dissimilarity (bottom row) across the two sampling areas.

### Environmental influence

The principal component analysis (PCA) allowed us to explore the relationship among the measured soil variables (see Table 1 S.I and Fig. 1 S.I.). The total inertia (the total variance) of the dataset was 43, with 35 unconstrained axes identified. The first principal component (PC1) had an eigenvalue of 14.50, explaining 33.72% of the total variance and represents an organic matter gradient. The second principal component (PC2) had an eigenvalue of 6.25, contributing 14.54% to the total variance and depicts a gradient of soil texture. Altogether, both PC1 and PC2 accounted for 48.26% of the variance.

Then, the main gradient in plot species composition was described for the first axis (Axis 1) of the NMDS ordination (Figure 3). Specifically, the Axis 1 identified a gradient of soil development, discriminating between plots with higher organic matter content and finer soil particle sizes (positive values) and those characterized by lower organic matter content and coarser soil particles (negative values). Furthermore, plots in burned area were grouped around the central area of the NMDS plot, without a clear pattern of distinction among them, but presenting a higher dominance of *Halimium calycinum* e *Corema album* in the coastal zone plots. On the other hand, the unburned area plots present a more prominent division along the gradient, with coastal and intermediate zone plots presenting poor soils dominated by *C. album* and *Stauracanthus genistoides* while the interior zone plots present richer soils dominated for species such as *Daphne gnidium, Lavandula angustifolia e Salvia rosmarinus.* Regarding species, three groups can be noticed across Axis 1: A first one, comprising *C. album* and *S. genistoides* both which dominate the coastal plots of both sampling areas, as well as in the intermediate plots of the unburned area; *H. calycinum*, *Cistus salviifolius* and *Halimium halimifolium* form a second group located near the center of the first NMDS; The remaining seven species (*Osyris lanceolata*, *S. Rosmarinus*, *Cytisus grandifloras*, *Daphne gnidium*, *L. angustifolia*, *Pinus pinea*, and *Juniperus phoenicea*) form a third group, primarily corresponding to unburned interior plots with more developed soils.

**Figure 3 –.**
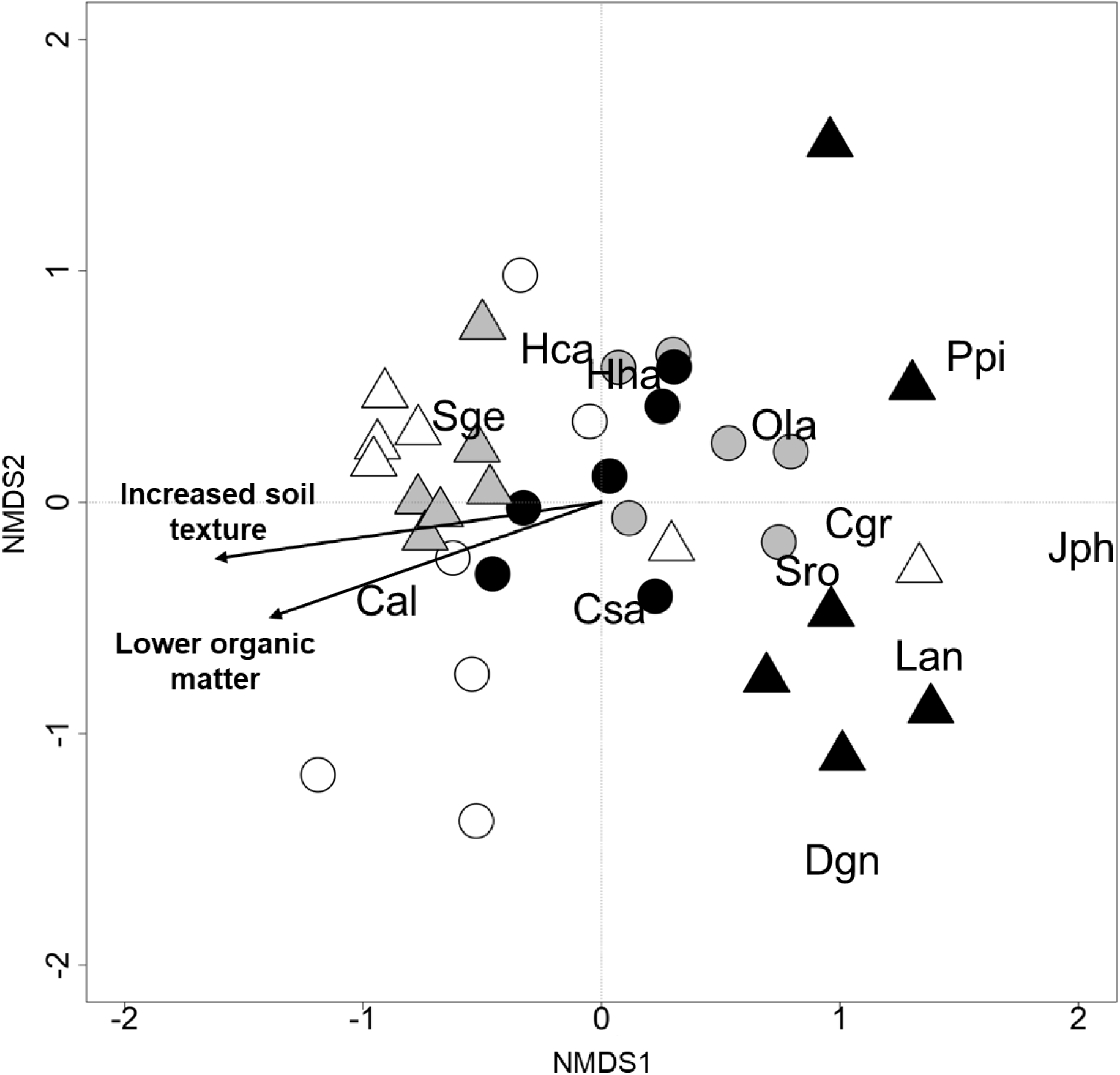
Axes 1 and 2 of the two-dimensional Non-Metric Multidimensional Scaling (NMDS) of sampling plots based on plant species cover (final stress 16.5 %). Circles represent plots in burned areas and triangles plots in unburned areas. Colours reflect location: white for coastal zone plots, grey for intermediate zone plots, and black for interior plots. Arrows indicate the main environmental gradients identified through correlation analysis.

### Indicator Species Analysis

The indicator species analysis revealed significant associations between certain plant species and specific groups based on both area and zone (Table 2 S.I.). When grouped by area, *Halimium halimifolium* was strongly associated with the burned area, showing an IndVal statistic of 0.898 and a highly significant p-value of 0.001. Additionally, *Lavandula angustifolia* was also significantly associated with the unburned area, with an IndVal statistic of 0.527 and a p-value of 0.046. In the analysis grouped by zone, *Lavandula angustifolia* also demonstrated a significant association with the interior transect, with an IndVal statistic of 0.563 and a p-value of 0.024.

### Species richness and abundance

The results of the analysis of variance, which evaluate the effects of area, transect, and their interaction on the abundance of woody plants, are presented in Table 2 and Fig. 4. The results show a significant effect of fire on woody plant abundance (F = 16.53, p < 0.001), with the burned area having higher abundance than the unburned area across all transects, while the differences between zones were smaller and only marginally significant, with a slight trend for higher abundance in the intermediate transects, especially in the burned area.

**Figure 4 –.**
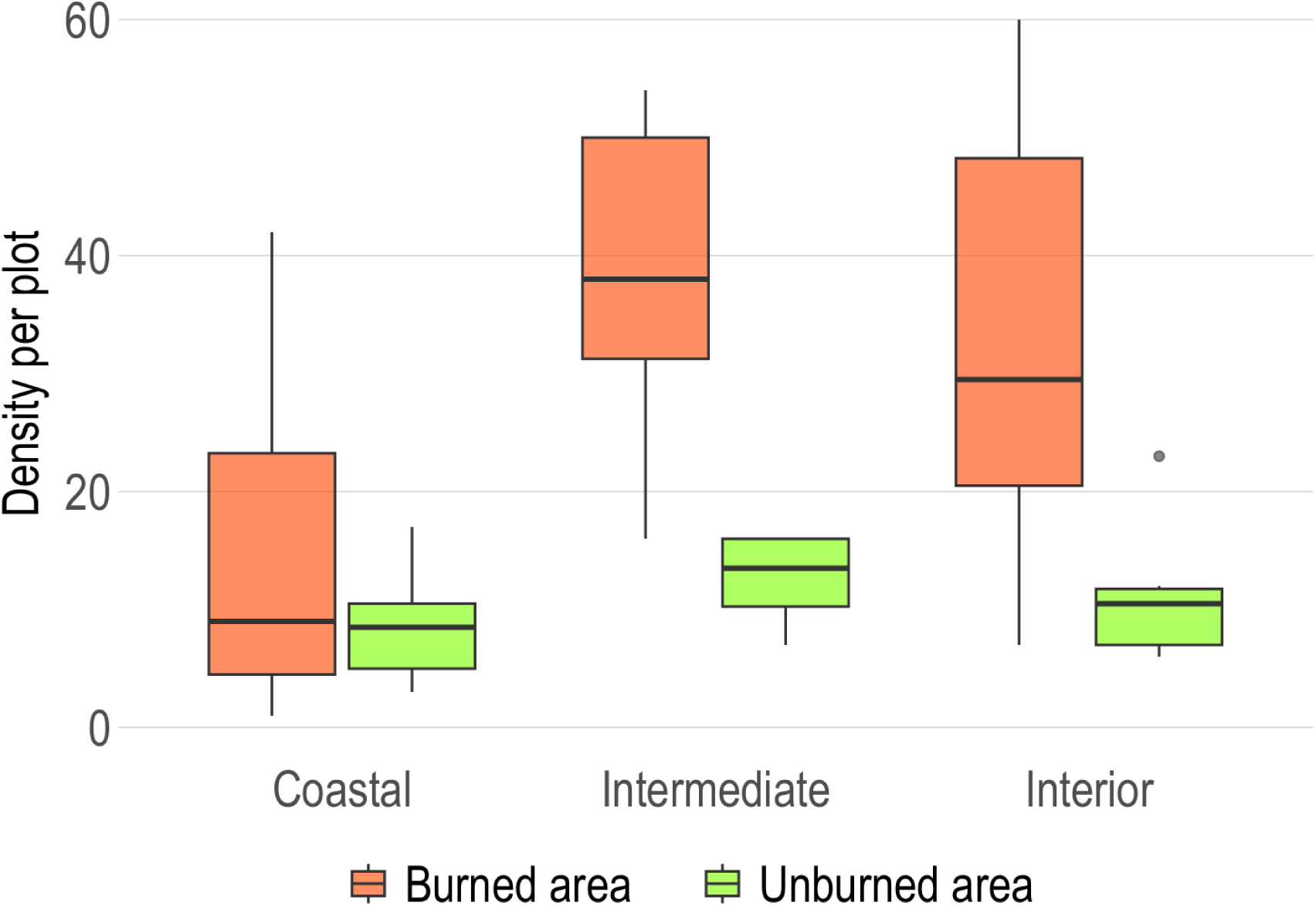
Number of woody plant species individuals per plot across coastal, intermediate, and interior zones in burned and unburned areas

**Table 2 –.**
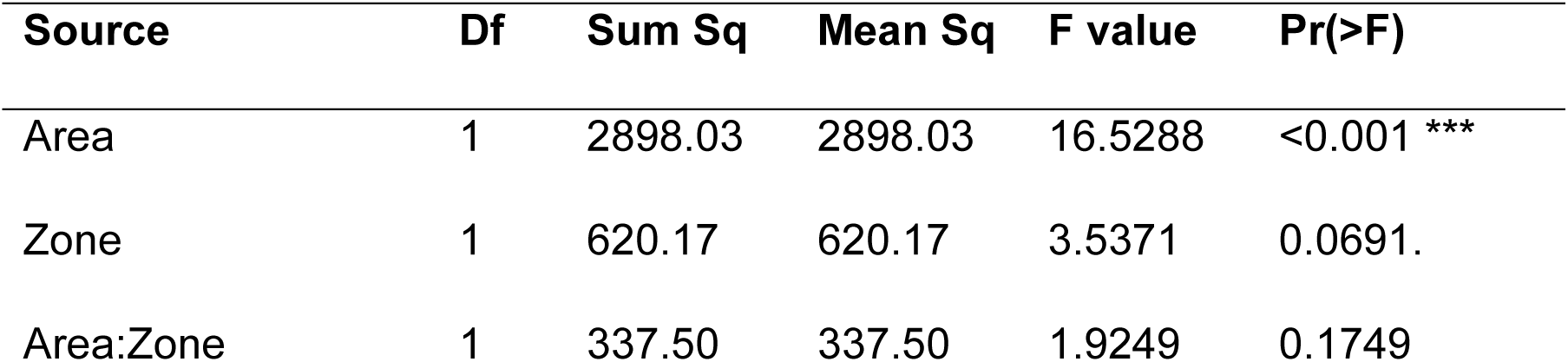
Table containing the results of the Anova for the woody plant abundance linear model.

The analysis of variance comparing species richness across area (e.g. the effect of fire) and zones (see Fig. 5 and Table 3) indicate a significant effect of zone on woody species richness (χ^2^ = 13.5, p = 0.01). In contrast, no significant effect of fire or the interaction between area and zone on species richness was detected.

**Figure 5 –.**
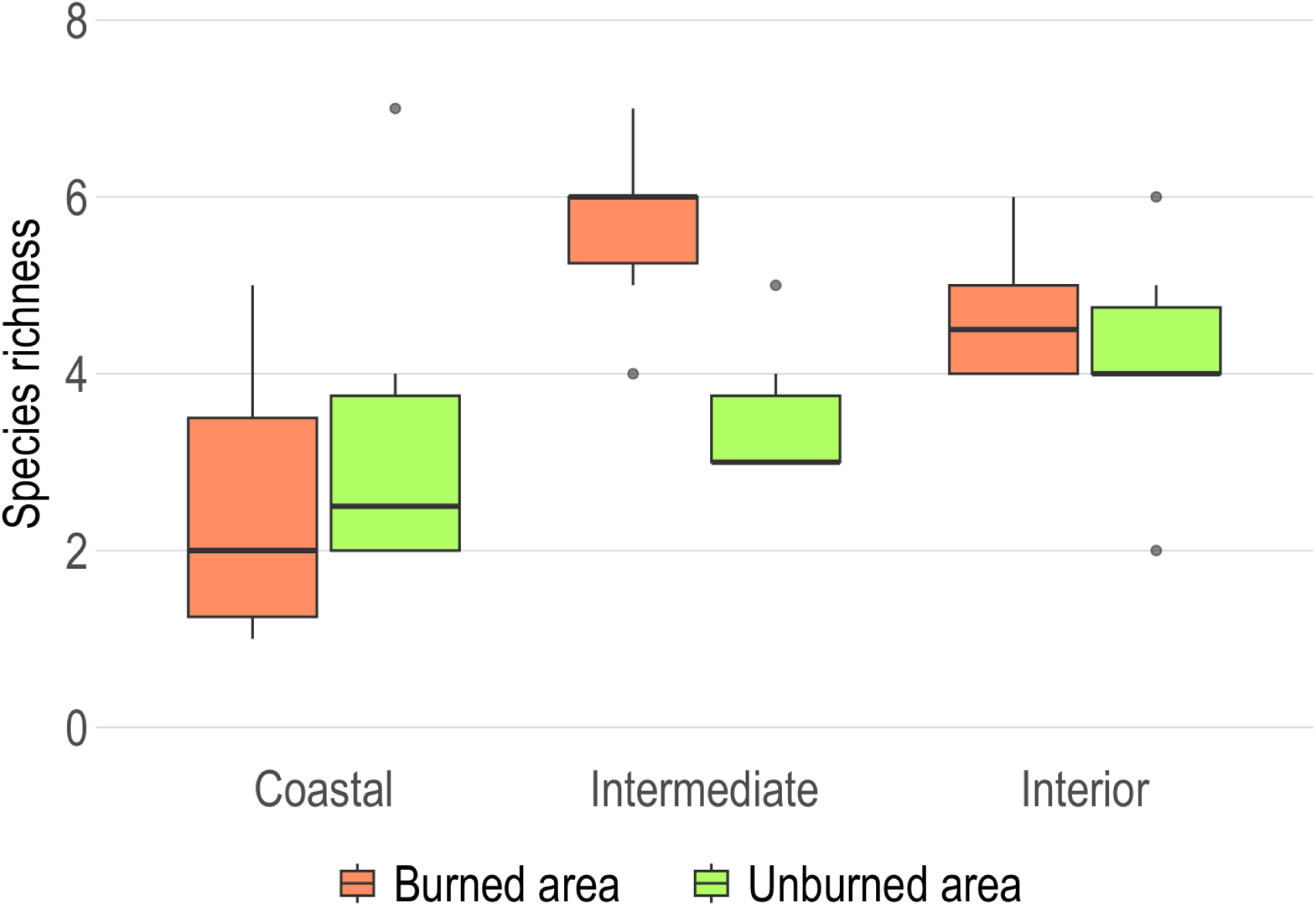
Comparison of the woody plant species richness present in the three samples transects for each area.

**Table 3 –.** Results of the Poisson GLM model assessing the effects of area, zone, and their interaction on woody species richness.

| Source | Df | Deviance | Residual Df | Residual Dev | Pr(>Chi) |
| --- | --- | --- | --- | --- | --- |
| Area | 1 | 3.361 | 34 | 87.611 | 0.2198 |
| Zone | 1 | 13.500 | 33 | 74.111 | 0.0139 * |
| Area:Zone | 1 | 2.667 | 32 | 71.444 | 0.2744 |

### Co-occurrence patterns

At the burned area, the observed C-score (6.54) fell within the 95% confidence interval of the simulated scores (mean = 6.34, 95% CI = 5.86 to 7.04, SES = 0.64), suggesting random co-occurrence patterns among woody species (Fig. 6 upper histogram). In contrast, at the unburned area, the observed C-score (9.83) exceeded both the mean simulated score (0.07) and the upper 95% confidence interval (8.71 to 9.56, SES = 3.56), indicating non-random co-occurrence with repulsion being the dominant pattern among species (Fig. 6 bottom histogram).

**Figure 6 –.**
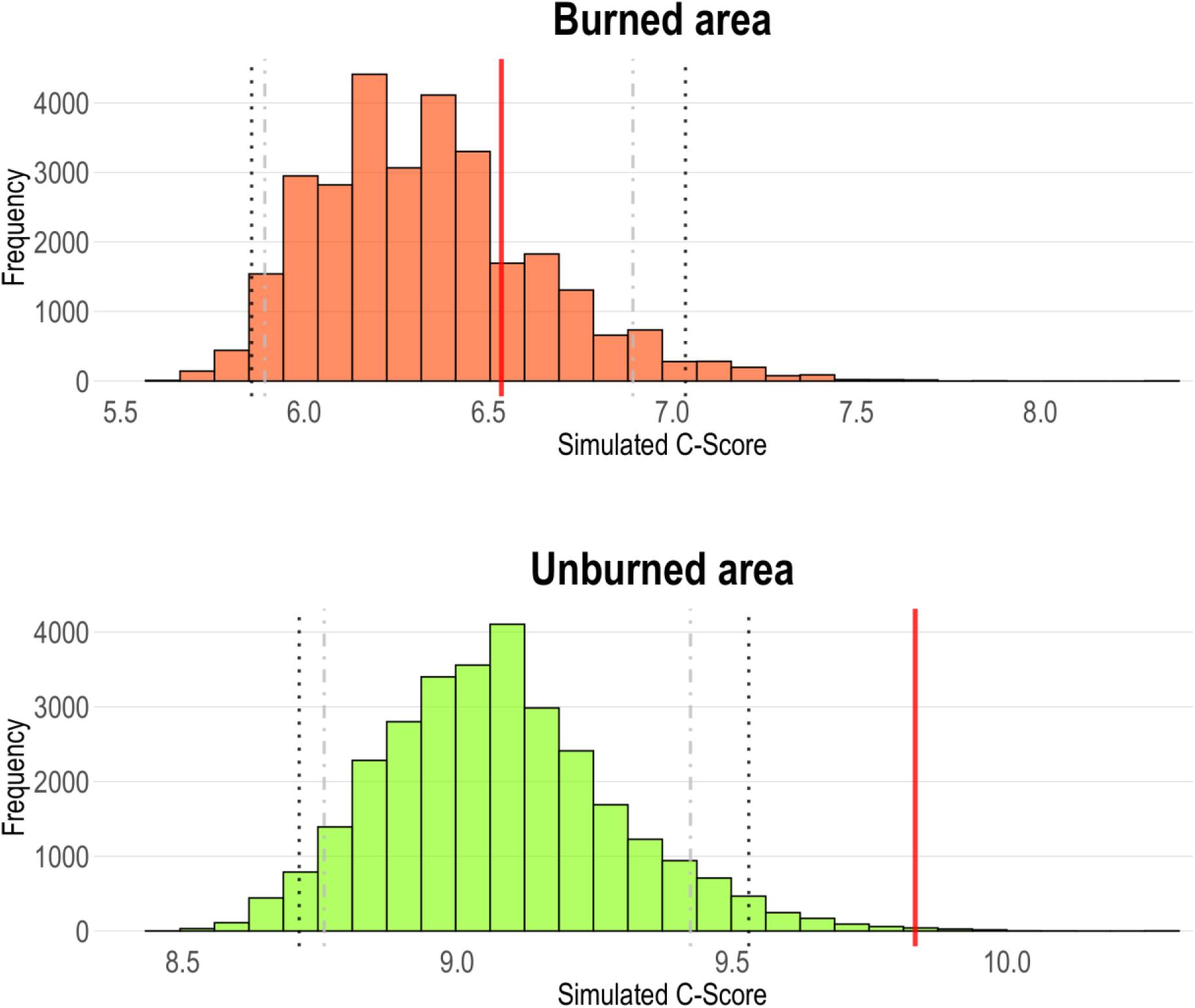
Histograms showing the distribution of simulated C-scores for the burned area (top) and unburned area (bottom). The red vertical line represents the observed C-score for each area. Dashed vertical lines indicate key simulated metrics, with black dashed lines marking the lower and upper 95% confidence intervals (two-tailed, grey – 1-tail, and black – 2-tail).

### Spatial structure of the community

The modularity analysis done to the 11 woody species sampled in the burned and unburned areas revealed notable differences in the woody plant community structure (Fig. 7). The results of this analysis were consistent for each sampling area throughout the three transects. For instance, in the burned area, all networks indicate the absence of internal species clustering, and present modularity values of 0 (or near 0, in the case of the intermediate transect). However, in the unburned area, all networks present higher levels of modularity, suggesting a more defined community structure.

**Figure 7 –.**
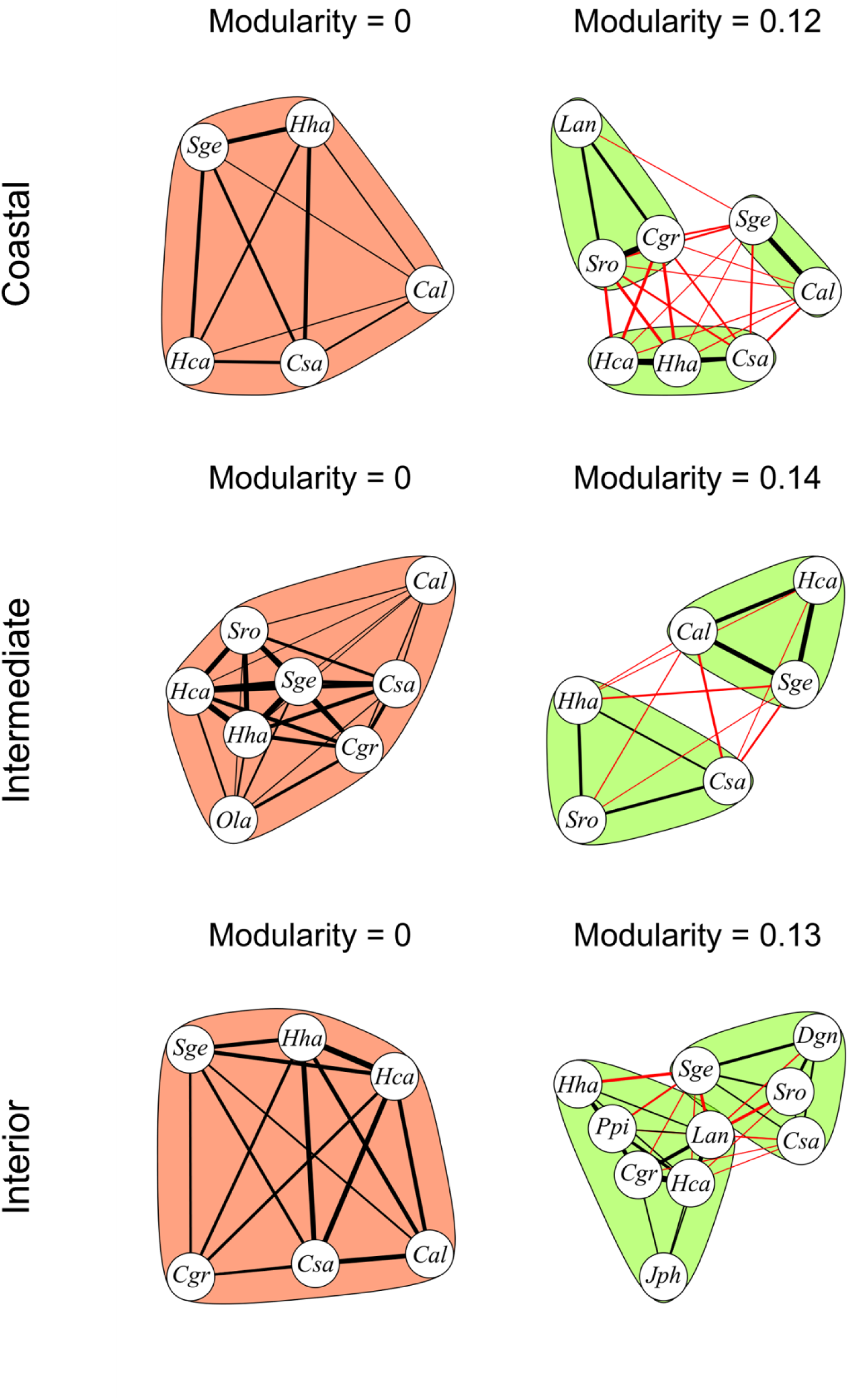
Image containing network diagrams of the woody plant species communities across the three sampling zones in the burned area (orange) and the unburned area (green). The network diagrams include nodes (which represent species) connected by lines which represent their interaction (being present in the same plot). The thickness of the lines indicates the frequency of interaction and the arrangement of species within each outlined colored section illustrates their modular grouping as determined by the Louvain method.

### Spatial Pattern Analysis

#### a) Generalized spatial patterns of woody plant community

To examine the overall associations of the woody plant community, a binomial logistic regression model was applied to each distance interval. This approach aimed to assess how fire influenced the overall pattern of spatial associations among woody plant species across the sampling areas (Fig. 8). The burned area exhibited a significantly higher number of positive spatial associations at the smallest distance interval (Distance I) when compared to the unburned area (coef = −2.41, p < 0.001). However, no significant differences were observed between the burned and unburned areas at the medium (Distance II) or largest (Distance III) intervals. Within the burned area, spatial associations varied across zones at the medium and largest distance intervals, with the interior zone showing more positive spatial associations than the coastal transect (Distance II: coef = 1, p = 0.014; Distance III: coef = 1.19, p = 0.009). Moreover, in the unburned area, both the intermediate and interior zones exhibited significantly more positive spatial associations when compared to the coastal transect. Specifically, the intermediate zone showed a stronger effect (coef = 2.22, p < 0.001), followed by the interior zone (coef = 1.59, p < 0.01).

**Figure 8 –.**
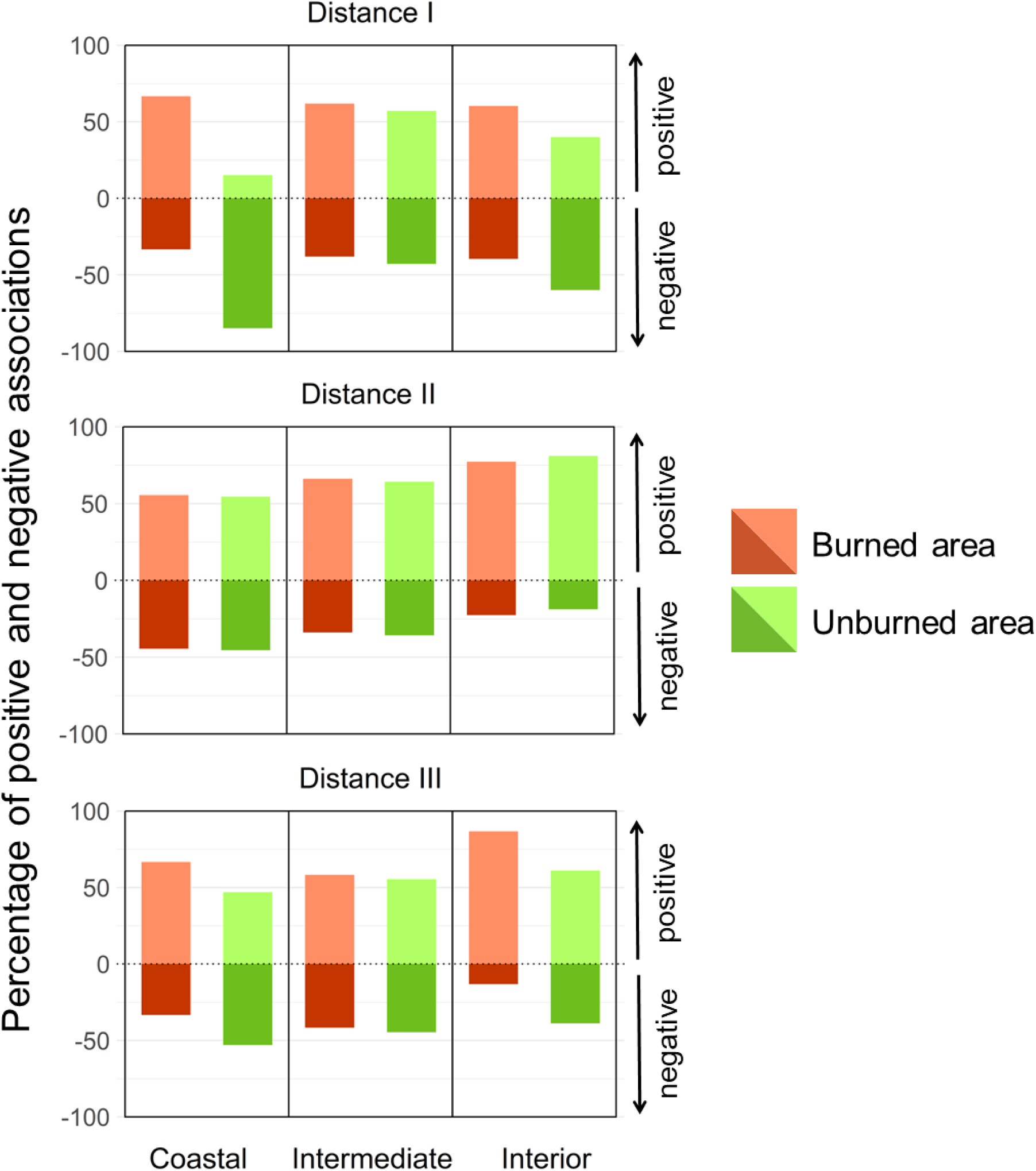
Figure representing the proportion of positive and negative interactions detected at three different ranges across the three transects in the two sampling areas.

#### b) Spatial patterns of recurrent woody plant species

Six species were identified in both sampling areas: C. album, *C. grandiflorus*, *H. calycinum*, *H. halimifolium*, *S. rosmarinus*, and *S. genistoides* (Fig. 9). For *C. album*, the number of detected positive and negative spatial associations did not differ significantly with fire or distance interval. However, a significant interaction was observed between fire and distance interval (χ^2^ = 112.6, p = 0.01) as well as between spatial associations and distance interval (χ^2^ = 118.8, p = 0.04). *C. grandiflorus* demonstrated an overall significant difference in the number of positive and negative spatial associations (χ^2^ = 197.62, p = 0.03), along with a significant difference between burned and unburned areas (χ^2^ = 191.21, p = 0.01). Additionally, a significant interaction was detected between the type of spatial association and distance interval (χ^2^ = 171.74, p < 0.001). *H. calycinum* exhibited significant differences in the number of positive and negative spatial associations (χ^2^ = 109.02, p < 0.001) and between burned and unburned areas (χ^2^ = 84.06, p < 0.001). Furthermore, a significant interaction was found between the type of spatial association and distance interval (χ^2^ = 77.80, p = 0.01). Similarly, *H. halimifolium* showed significant differences in the number of positive and negative spatial associations (χ^2^ = 152.70, p < 0.001) and between burned and unburned areas (χ^2^ = 81.63, p < 0.001). For *S. rosmarinus*, no significant effects were detected for any of the predictors. Finally, *S. genistoides* exhibited significant differences in the number of positive and negative spatial associations (χ^2^ = 86.71, p < 0.001) and between burned and unburned areas (χ^2^ = 80.15, p = 0.01).

**Figure 9 –.**
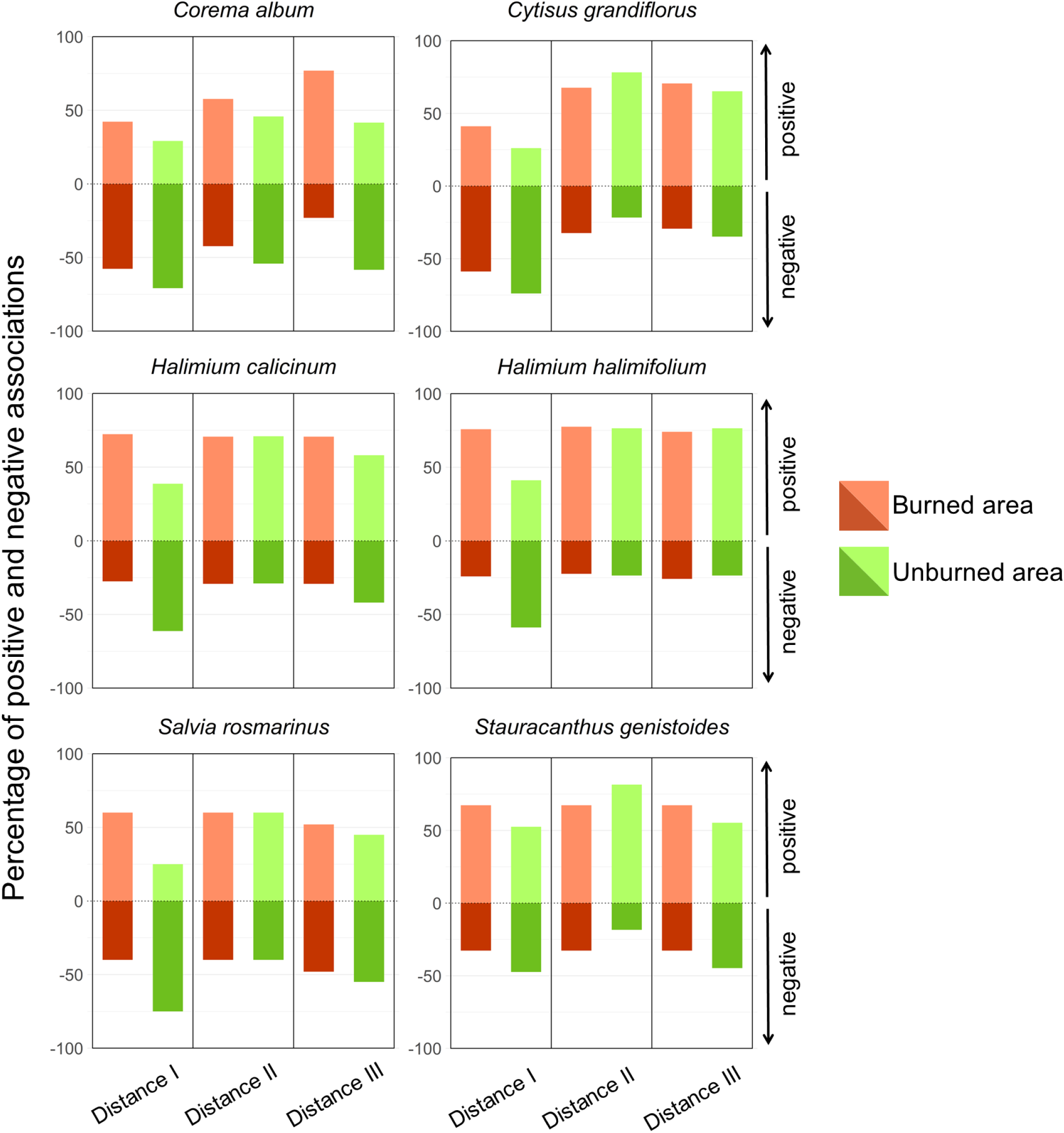
Percentage of positive and negative spatial associations for the six woody plant species found in both sampling areas (*Corema album*, Cytisus grandiflorus, Halimium calycinum, *Halimium halimifolium*, *Salvia rosmarinus*, and *Stauracanthus genistoides*) across three distance intervals (I, II, and III) in burned (orange) and unburned (green) areas. Positive associations are represented above the dotted line, while negative associations are shown below.

## Discussion

Overall, our results reveal how a disturbance caused by fire can affect the core dynamics of a community of woody plants. In the unburned area (Laguna del Jaral), species composition is strongly influenced by the coastal-to-inland gradient and soil characteristics, with well-structured communities shaped by deterministic processes ^42^. In contrast, the burned area (Cuesta Maneli) exhibits more homogeneous species composition across the gradient, with greater overall abundance but no differences in species richness. Species in the burned area co-occur randomly, suggesting neutral interactions to be more common, while unburned areas display structured patterns dominated by negative spatial associations, suggesting a stronger effect of competition. Furthermore, spatial analyses show that positive spatial associations occur more frequently in the burned area, particularly at smaller distance intervals. This finding reinforces the hypothesis of reduced competition following fire and highlights the critical role of interactions between nurse plants and facilitated species in driving vegetation dynamics within this system ^58^. Likewise, the six species present in both sampling areas showed a similar pattern, where four of them (*C. grandiflorus*, *H. calycinum*, *H. halimifolium*, and *S. genistoides*) showed a significant effect of fire in their spatial associations, all presenting higher percentage of positive spatial associations at the smaller distance intervals.

### Woody species community composition

Fire did not appear to affect overall community composition when comparing burned and unburned areas. However, it significantly disrupted the community composition along the coastal-inland gradient. This disruption likely stems from changes in soil properties caused by fire. In the burned area, woody plant species were less influenced by soil parameters, likely due to the homogenizing effects of fire. Fire is known to reduce variability in soil attributes by redistributing nutrients and altering microtopography ^59–61^, thereby diminishing the constraints typically imposed by soil conditions. This homogenization may explain the lack of correlation between soil dissimilarity and species composition in the burned area, reflected by the absence of clear grouping of plots in the burned area along the soil texture and organic matter gradients. These conditions, coupled with reduced competitive pressure, enabled *H. halimifolium* to flourish, making it an indicator species for the burned area. On the other hand, the unburned area exhibits stronger spatial sorting, which indicates a more heterogeneous community along the coastal-inland gradient. Notably, this environmental gradient appears to favor certain species, like *C.* album and *S. genistoides*, which demonstrate a shared resilience to similar environmental conditions ^7^. Such conclusions can also be supported when we take into consideration the fact that woody plant composition in the burned area does not vary among zones at the same rate as it does in unburned one. Furthermore, similar patterns can be inferred for *L. angustifolia*, which was identified as an indicator species for the unburned area. However, the strength of its association with the unburned area was weaker compared to the stronger and more pronounced link observed between *H. halimifolium* and the burned area. The unburned area, characterized by greater environmental heterogeneity, supports spatial variation in species presence, contributing to a less distinct and more variable species composition compared to the more homogenous burned area.

Species abundance was also notably impacted by the fire disturbance. Beyond its homogenizing effects, fire can enrich the nutrient content of the soil. This happens due to the production of ash following a fire, which leads to an increase of soil organic matter, pH, electrical conductivity and several elements essential to vegetation growth, such as carbon, calcium, magnesium, sodium and nitrogen ^61^. The soil nutrients vectors from the PCA analysis did not show a distinction between the sampling area, which suggests that soil has already recovered from the effects of the fire. However, it can be inferred that the nutrient enrichment and the ecological spaces created by fire contribute to a greater abundance of species, as reflected in the observed effects of fire. Additionally, fire disturbance is known to affect dune topography, as the absence of plant cover leaves sand banks exposed to eolian transport ^24,62^.

Considering these effects, we propose that fire has reduced the community complexity in the burned area. The 2017 fire disrupted the coastal-inland gradient, redistributed nutrients, and created new vacant ecological spaces. These changes allowed the succession in which the existing woody plant species exploit the conditions, with new sprouts and seedlings established in areas where such abundance was previously unattainable due to competitive pressures ^63^. These findings suggest several homogenizing effects of fire disturbance that may have diminished the natural gradient-driven effects across species succession. This has resulted in a more uniform community, likely in a transient state, which is expected to be reshaped by biotic interactions.

### Community structure

As the burned area seems to stand in a transitional state, neutral processes (such as stochastic dispersal or environmental variability that equally affects all species) seem to be the main responsible for the structure of the community. Additionally, the burned area exhibits a notable lack of internal clustering among woody plant species (null modular values), resulting in a diffuse community structure with weak or absent associations due to relatively uniform habitat conditions. Conversely, in the unburned area, competitive interactions appear to play a more significant role in structuring the community. The pronounced spatial and environmental gradients observed in the unburned area promote niche differentiation, leading to increased negative spatial associations among co-occurring species as detected by the C-score analysis. These interactions may account for the observed clustering of species. Additionally, higher modulation values indicate that woody plant species are forming distinct assemblages, potentially driven by environmental filtering and biotic interactions. These structural differences between the burned and unburned woody plant communities highlight the substantial impact of fire on this ecosystem. Regarding the impact of fire on spatial associations, at the smallest distance interval, the burned area exhibited significantly higher levels of positive spatial associations, which may suggest that facilitative interactions might be more prominent in this area (such as nursing). This pattern contrasts with the unburned area, particularly in the coastal transect, where lower levels of positive spatial associations were observed. Indeed, fire significantly altered species’ spatial associations, resulting in a higher number of positive spatial associations in the burned area compared to the unburned area. This contrasts with the unburned area, where competitive interactions appear to play a more dominant role in shaping the community structure. At this scale, post-fire conditions likely foster facilitative interactions among successional woody plant species, possibly due to diminished competition and enhanced resource availability, while in the unburned area, the woody plant community seems to be more tightly structured by competitive exclusion and niche differentiation. Furthermore, the detected differences in spatial associations across distances suggest that the effect of fire is scale dependent. As positive spatial associations are more evident at smaller scales, larger scales may convey a more complex interplay between facilitation and competition depending on the zone and area. Overall, considering the increase of positive associations and lack of strong competition, we highlight the transitional state of the woody plant community, where neutral or facilitative processes dominate, potentially leading to a gradual re-establishment of competitive dynamics over time as the woody community stabilizes.

Concerning the impact of fire in the spatial associations of recurrent plant species, we’ve detected species-specific impact, with varying effects across different distance intervals. For instance, *C. grandiflorus*, *H. calycinum*, *H. halimifolium*, and *S. genistoides* exhibit notable shifts in their spatial interactions in the burned area, indicating a strong disruption of their typical patterns. Furthermore, the interaction between the effect of fire and distance intervals also shows its relevance in shaping spatial associations, when species such as *C. album*, vary their spatial associations across scales. On the other hand, *S. rosmarinus* shows no significant changes, suggesting resilience to fire disturbance. Such findings go in line with what was previously mentioned, in which an increase in positive spatial associations in burned areas suggests a reduction in competitive pressures, leading to more facilitative interactions, while unburned areas show more distinct clustering due to stronger competition and environmental gradients. Ultimately, these findings suggest that fire homogenizes species associations in burned areas, while unburned areas maintain more differentiated community structures due to competitive dynamics.

### Woody plant succession

Finally, our results are in line with Egler’s Initial Floristic Composition hypothesis ^64^. According to this hypothesis, the initial species composition has a strong influence on the trajectory of community development. This effect is evident in the similarity of species richness between both areas. Furthermore, our results suggest that in the burned area, currently negative interactions between species are not a primary driver of community structure, implying that species are passing through the abiotic filter and benefiting from positive interactions, such as facilitation. The differences in species abundance between the sampling areas suggest that the woody plant community in the burned area is still in the process of development. Our observations provide a “snapshot” of this evolving community, which already exhibits the same species richness of a nearby locality but is likely continuing to undergo shifts in composition and structure, as negative biotic interactions become more relevant. Indeed, these observations contrast with the Clement’s Relay Floristics model ^65^, which proposes that plant succession is a stage-based process where early colonizers do not necessarily influence the final community composition. Indeed, it has already been suggested that ecological communities are not cohesive entities nor follow a predictable successional stage. They are shaped by the individualist response of species to many variables, factors, chance and other species, which gives them a more fluid and dynamic structure, which was also hypothesized by Gleason 1926 ^66^. Numerous studies have already discussed the limitations of Clements’ work ^9,67^, and our results further demonstrate that succession does not always follow a predictable, stage-driven pattern.

## Conclusion

This study shows that fire significantly impacts woody plant communities in coastal regions by homogenizing species composition and shifting community assembly processes. In the unburned area, species composition is structured by the coastal-to-inland gradient and competitive interactions, resulting in heterogeneous communities. In contrast, the burned area exhibits random species co-occurrence, indicating neutral and facilitative interactions dominate post-fire. Moreover, we identified that fire reduces competitive pressures, increasing positive spatial associations. The burned community is currently transitional state likely giving way to competitive dynamics over time. These findings challenge traditional succession models, emphasizing the dynamic nature of community assembly influenced by both original species composition and post-disturbance interactions.

## Supplementary Information

**Table 1 S.I. -.**
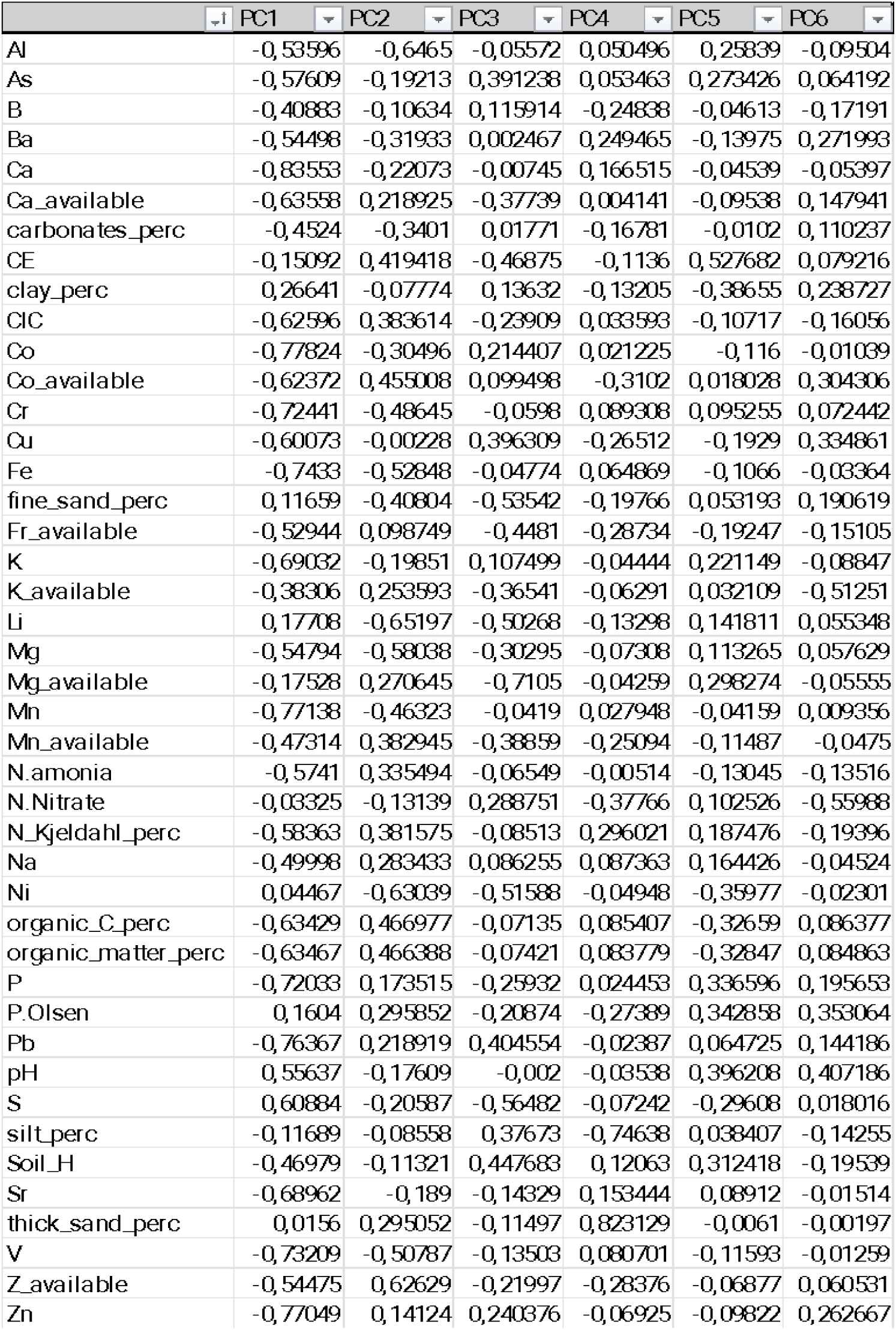
Species scores (shown for the first six principal components) are provided for each variable. These scores represent the projections of the onto the principal components.

**Figure 1 S.I. –.**
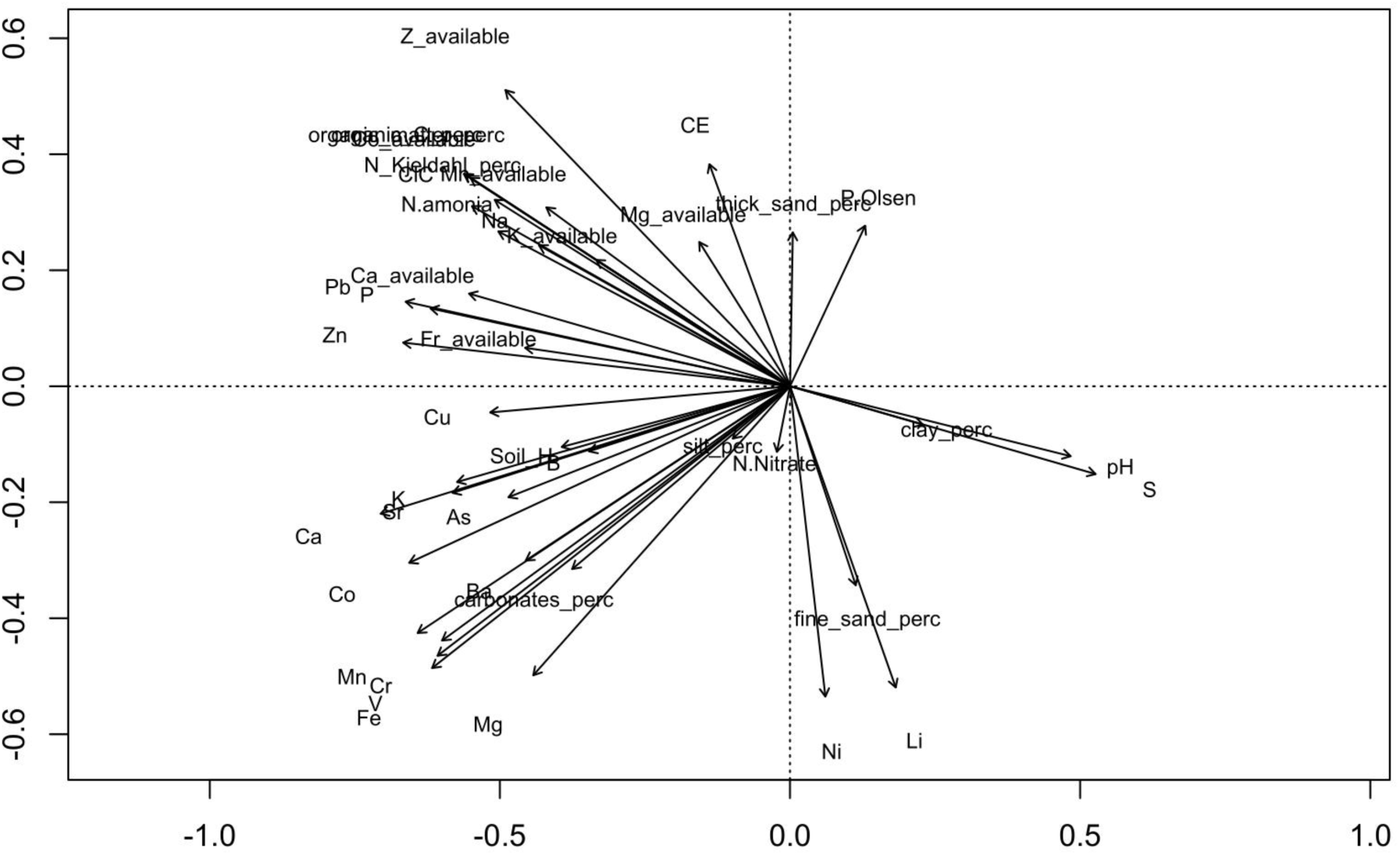
Figure containing a visual projection of the used variables onto the two first principal components.

**Table 2 S.I. -.** This table presents the results of the Indicator Species Analysis for the plant species dataset. The analysis identifies species that are significantly associated with either Location or Zone.

|  | Species | Area | IndVal<br>Statistic | p-value |  |
| --- | --- | --- | --- | --- | --- |
| Indicator Species by area | *Halimium halimifolium* | Burned | 0.898 | 0.001 | *** |
|  | *Lavandula angustifolia* | Unburned | 0.527 | 0.046 | * |
| Indicator Species by zone | Species | Zone | IndVal<br>Statistic | p-value |  |
|  | *Lavandula angustifolia* | Interior | 0.563 | 0.024 | * |

